# On the need of individually optimizing temporal interference stimulation of human brains due to inter-individual variability

**DOI:** 10.1101/2025.01.13.632831

**Authors:** Tapasi Brahma, Alexander Guillen, Jeffrey Moreno, Abhishek Datta, Yu Huang

## Abstract

**Introduction:** Transcranial temporal interference stimulation (TI, TIS, or tTIS), also known as interferential stimulation (IFS), is able to focally stimulate deep brain regions, provided it is properly optimized. We previously presented an algorithm for optimizing TI using two arrays of electrodes and showed that it can achieve more focal stimulation compared to optimized high-definition transcranial electrical stimulation (HD-TES) and conventional optimized TI using two pairs of electrodes, especially in the deep brain areas such as the hippocampus. However, those modeling studies were only performed on an averaged head (MNI152 template) and three individual heads without exploring inter-individual variability. Existing TI works in the literature mostly utilize a common (possibly optimized) montage of two pairs of electrodes on different individual heads without considering inter-individual variability.

**Material and method:** Here we aim to study the inter-individual variability of optimized TI by applying the same optimization algorithms on N = 25 heads using their individualized head models. Specifically, we compared the focality achieved by different stimulation techniques at six different regions of interest (ROI; right hippocampus, left dorsolateral prefrontal cortex, left motor cortex, right amygdala, right caudate, and left thalamus) under both individually optimized and unoptimized montages. We also conducted numerical sensitivity analysis on the individual optimization and performed phantom recordings to test our models.

**Results:** As expected, there is a variability in focality achieved by TI of up to 1.2 cm at the same ROI across subjects due to inter-individual differences in the head anatomy and tissue conductivity. We show that optimized TI using two arrays of electrodes achieves higher focality than that from optimized HD-TES at the same level of modulation intensity at 5 of the 6 ROIs. Compared to using a common montage either optimized from the MNI152 template or from the literature, individually optimized TI using two pairs of electrodes improves the focality by up to 4.4 cm, and by up to 1.1 cm if using two arrays of electrodes. Focality achieved by the individual optimization is sensitive to random changes and can vary up to 9.3 cm due to the non-lienarity of TI physics. Experimental recordings on a head phantom confirms the drop in TI stimulation strength when using unoptimized montages as predicted by our *in silico* models.

**Conclusion:** This work demonstrates the need of individually optimizing TI to target deep brain areas, and advocates against using a common head model and montage for TI modeling and experimental studies.

## Introduction

Transcranial electrical stimulation (TES) delivers weak electric current (≤ 2 mA) to the scalp in order to modulate neural activity of the brain (Nitsche and Paulus, 2000). TES has been shown to improve cognitive functions and help treat some neurological diseases such as major depression (Bikson et al., 2008), epilepsy (Auvichayapat et al., 2013; Fregni et al., 2006b), Parkinson’s disease (Fregni et al., 2006a), chronic pain (Fregni et al., 2007), and stroke (Meinzer et al., 2016). Conventional TES using two sponge electrodes is shown to generate diffused electric fields and cannot reach deep targets in the brain (Datta et al., 2009). High-definition TES (Datta et al., 2009; Edwards et al., 2013) utilizing several small disc electrodes has thus been proposed to achieve more focal or more intense stimulation. Several algorithms have been developed along this line for optimizing HD-TES in terms of focality or intensity of the induced electric field (Dmochowski et al., 2011; Guler et al., 2016; Huang et al., 2018; Ruffini et al., 2014; Sadleir et al., 2012). However, even with optimized stimulation montages, modeling studies have shown that HD-TES is not able to focally stimulate deep brain regions (Dmochowski et al., 2011; Huang and Parra, 2019).

Recently transcranial temporal interference stimulation (TI, TIS, or tTIS), also known as interferential stimulation (IFS), has drawn much attention in the TES community as both computational and experimental studies show that TI can reach deep brain areas (Grossman et al., 2017; Huang et al., 2020; Huang and Datta, 2021; Violante et al., 2023). TI applies sinusoidal waveforms of similar frequency through two pairs of electrodes. The resulting interfered electric field is thus amplitude modulated containing both a fast oscillating carrier signal in the kHz range and a slowly oscillating modulation envelope in the low frequency range. The premise of TI is that neurons only respond to the slower oscillation due to their property of low-pass filtering (Grossman et al., 2017). Computational researchers have since been working on optimizing the placement of the two pairs of electrodes on the scalp to maximize stimulation focality. Most of these efforts are considering only two pairs of electrodes as originally proposed by Grossman et al (Lee et al., 2020; Rampersad et al., 2019; von Conta et al., 2021). Some are trying to extend to multiple (>2) pairs of electrodes but the underlying algorithms are either *ad hoc* (Lee et al., 2022), or not systematically optimizing the electrode locations (Xiao et al., 2019) or the full physics of TI (Cao and Grover, 2020). To the best of our knowledge, we are the first to formulate a rigorous mathematical framework to optimize the full physics of the interfered envelope of TI based on two arrays of multi-channel electrodes (up to 93 per frequency) instead of two pairs or multi pairs of electrodes (Huang et al., 2020). We showed *in silico* that compared to HD-TES, optimized multi-channel TI stimulates deep brain areas with much higher focality at the same modulation strength (Huang et al., 2020). We further showed that compared to optimized TI using two pairs of electrodes, optimized two-array TI also shows higher focality at the same modulation strength (Huang and Datta, 2021).

However, we only tested our optimization algorithms for TI on the averaged standard head (Huang et al., 2020) and up to three individual heads (Huang and Datta, 2021). Considering inter-individual variability in head anatomy (Datta et al., 2012) and tissue conductivity (Huang et al., 2017), we expect to see differences across subjects in the optimal montages for stimulating the same target location in the brain. Existing computational and experimental work in the literature mostly use a common head model or electrode montage to study TI (Grossman et al., 2017; Vassiliadis et al., 2024b; Violante et al., 2023). This may explain why TI did not show any effect when applied on the visual cortex of 10 subjects in a recent work (Iszak et al., 2023). In this paper we show that (1) optimal TI montages vary significantly across subjects; (2) optimized two-array TI shows higher focality compared to optimized HD-TES at most targets; (3) a common or a literature montage always gives worse focality under two-pair TI stimulation compared to individually optimized ones; (4) random TI montages further decreases the focality due to the non-linearity in TI physics; (5) recordings from a head phantom confirm the importance of individual optimization of TI montages. We therefore argue that individualized modeling and optimization is important in future TI studies.

## Materials and Methods

### Individualized head models

Ten individual T1-weighted magnetic resonance images (MRI) were randomly selected from the 1200 Subjects Data Release in the Human Connectome Project (Van Essen et al., 2012). As this database consists of young adults only (ages 22–35), to make our results more generalizable, we also included fifteen subjects from the Neurodevelopment database (Richards and Xie, 2015) with age spans from 17 to 89 years. In total 25 heads were modeled. Open-source toolbox ROAST (Huang et al., 2019) was used on each of these subjects to generate individualized lead-field that is needed for optimized HD-TES (Dmochowski et al., 2011) and optimized TI (Huang et al., 2020; Huang and Datta, 2021). ROAST generated the lead field with a simple one-line command (Huang, 2019). Specifically, the individual head MRI was segmented into gray matter, white matter, cerebrospinal fluid (CSF), skull, scalp, air cavities by the New Segment toolbox (Ashburner and Friston, 2005) in Statistical Parametric Mapping 12 (SPM12, Wellcome Trust Centre for Neuroimaging, London, UK) implemented in Matlab. Segmentation errors such as discontinuities in CSF and noisy voxels were corrected by a customized Matlab script (Huang et al., 2013). Since TES modeling work has demonstrated the need to include the entire head down to the neck for realistic current flow, in particular in deep-brain areas and the brainstem (Huang et al., 2013), the field of view (FOV) of the head MRI was extended down to the neck by zero-padding 30 slices when calling ROAST, which automatically pastes the lower part of the standard head published in Huang et al., 2013 to the voxel space of the head MRI. 72 high-definition disc electrodes following the convention of the standard 10/10 international system (Klem et al., 1999) were placed on the scalp surface by custom Matlab script (Huang et al., 2013). A finite element model (FEM, Logan, 2007) was generated from the segmentation data by open-source Matlab toolbox iso2mesh (Fang and Boas, 2009). A typical head model consists of about 700,000 nodes and 4 millions elements. Laplace’s equation was then solved (Griffiths, 1999) in getDP (Dular et al., 1998) for the electric field distribution in the head. Default tissue conductivities in ROAST were used (in S/m): gray matter – 0.276; white matter – 0.126; CSF – 1.65; bone – 0.01; skin – 0.465; air – 2.5×10^-14^; gel – 0.3; electrode – 5.9×10^7^. With one fixed reference electrode Iz as cathode, the electric field was solved for all other 71 electrodes with unit current density injected for each of them, giving 71 solutions for electric field distribution representing the lead field of each individual head. The overall modeling pipeline along with the optimization process is shown in Fig. 1. (Fig. 1 to be here)

**Fig. 1:**
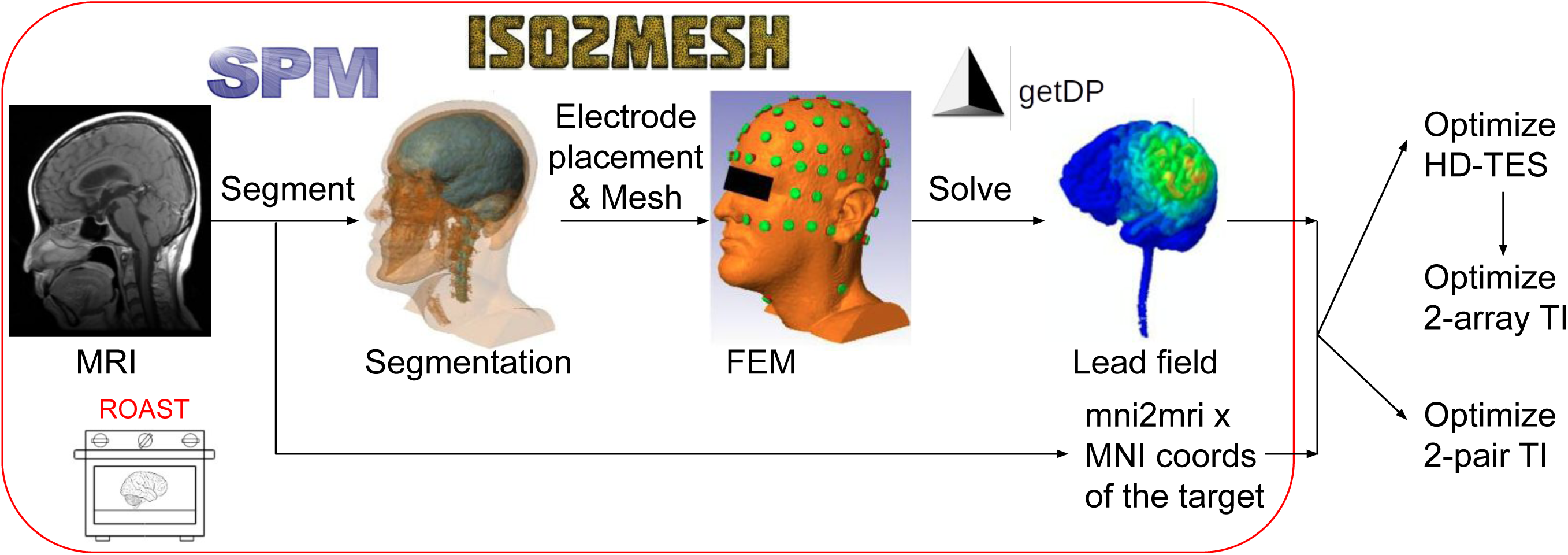
The modeling pipeline of ROAST for making individualized models of TES and TI (in the red round-rectangular), and the process of optimizing TES and TI. SPM: Statistical Parametric Mapping; MRI: magnetic resonance images; FEM: finite element model; MNI: Montreal Neurological Institute, a research institute that made the MNI space for brain mapping; mni2mri: a transform estimated by SPM to map the MNI coordinates (coords) of a target into the native MRI space.

### Optimization of HD-TES and TI

We used existing algorithms (Dmochowski et al., 2011; Huang et al., 2020; Huang and Datta, 2021) for optimizing HD-TES and TI to stimulate 6 target locations in the brain: right hippocampus (MNI coordinates [28,-22,-14]), left dorsolateral prefrontal cortex (DLPFC, MNI coordinates [-39,34,37]), left motor cortex (MNI coordinates [-48,-8,50]), right amygdala (MNI coordinates [21,-1,-22]), right caudate (MNI coordinates [14,13,11]), and left thalamus (MNI coordinates [-9,-17,6]). All these target locations except the DLPFC and motor cortex are in deep brain regions, as we want to fully leverage the advantage of TI in reaching the deep brain as shown in previous modeling and experimental studies (Grossman et al., 2017; Huang et al., 2020; Violante et al., 2023). The MNI coordinates were obtained using an online tool available at https://bioimagesuiteweb.github.io/webapp/mni2tal.html. These MNI coordinates were mapped into each individual head using the transform estimated by the segmentation algorithm utilized by ROAST (Ashburner and Friston, 2005; Huang et al., 2019; Fig. 1). We solved the optimization for 2-array TI and HD-TES at the same time. Specifically, we initialized the optimization for 2-array TI using solutions from max-focality HD-TES at the same target location. Optimizations of both 2-array TI and HD-TES aim to maximize the stimulation intensity at the target while limiting the stimulation energy in the non-target area. For HD-TES, this could be implemented by the classic least-constraint-minimal-variance (LCMV) algorithm (Dmochowski et al., 2011). For TI, this optimization problem is non-convex as the stimulation intensity (i.e., modulation depth, see below) is a non-linear and non-convex function of the injected current on the scalp. We solved this by sequential quadratic programming (Brayton et al., 1979; Huang et al., 2020). As shown in Fernández-Corazza et al., 2020, the optimization goal intrinsically encodes a trade-off between the focality and intensity of the modulation at the target, and this trade-off could be modeled by the energy constraint at the non-target areas. Therefore, we implemented the optimization by varying the energy constraints at non-target areas for 12 different levels (Huang et al., 2020), each time initializing the 2-array TI optimization with solutions from HD-TES optimization. For safety consideration, the total injected current was limited to 2 mA. For 2-pair TI optimization, we performed an exhaustive search to find the best two pairs of electrodes that achieves the highest focality of modulation at the target. Here focality was defined as the size of the brain volume stimulated at 50% (or more) of the modulation at the target (similar to the concept of full width at half maximum (FWHM) in classic imaging (Gonzalez and Woods, 2002)). Specifically, the search was performed across: (1) four possible electrodes selected from the 72 placed electrodes on the scalp; (2) three possible ways of pairing in these four electrodes for the two frequencies used in TI; (3) amplitudes of injected current at one pair of electrodes, in the range of 0.5 mA to 1.5 mA, with a step size of 0.05 mA and the sum across the two pairs of electrodes being 2 mA. In total 64,813,770 montages were considered for one target in the head model (1,028,790 possible combinations of 4 electrodes × 3 possible ways of pairing × 21 amplitudes of injected current, Huang and Datta, 2021). Note for all optimization problems, we optimized the modulation depth along the radial direction, i.e., the direction pointing from the target location to the center of the brain (defined as MNI coordinates of [0,0,0]). The modulation depth is the strength of the envelope of the interfered signal from the two pairs (or arrays) of electrodes. For more technical details, one is referred to Huang, 2023; Huang et al., 2020; Huang and Datta, 2021; Huang and Parra, 2019.

### Comparison between optimized HD-TES and optimized TI

Different levels of energy constraints at the non-target areas give different optimal electrode montages, as represented by a focality-modulation curve (Huang et al., 2020). Solutions under very strict (i.e., small) energy constraints usually result in electrode montages that do not use up all the 2 mA budget of injected current (Huang et al., 2020). We therefore discarded these solutions when analyzing the results. When showing examples of electrode montages across subjects and stimulation modalities (HD-TES and 2-array TI), we used the strictest energy constraint at which the injected current just reaches the total budget of 2 mA, i.e., the left-most point on the focality-modulation curve. Note that at the same level of energy constraint, HD-TES and 2-array TI may not return the same modulation depth. To facilitate comparison of focalities between these two stimulation modalities, we made sure that HD-TES outputs the same modulation at the target as that from 2-array TI by fine-tuning the energy constraint for HD-TES. As we will see in the Results section, the focality-modulation curve spans for different subjects at different locations in the focality-modulation space. To facilitate comparison across subjects and stimulation modalities, we mapped the curve for each subject to a normalized modulation depth between 0% and 100% by linear interpolation. Note for 2-pair TI, no energy constraint was explicitly specified in the searching algorithm, so the optimal montage is unique for a specific target on a specific subject.

### Comparison of optimized vs. common and literature montages

To show the performance of using a common montage on individual heads compared to that of using individually optimized montages, we also computed the optimal montages of HD-TES, 2-array TI, and 2-pair TI on the MNI152 head (Grabner et al., 2006). We then applied these optimal montages from the MNI152 head (i.e., common montages) onto all the 25 individual heads, computed the achieved stimulation focality at the 6 targets, and compared them to those from using individually optimized montages. Note that for HD-TES and 2-array TI, we again calculated the results for 12 different levels of energy constraints, and performed the same normalization process on the modulation depth to facilitate the comparison. For 2-pair TI there is only one unique solution for a specific target on a specific head.

As 2-pair TI is more popular than 2-array TI in the TI literature, we also searched for the montages used in the TI literature to target the 6 targets, and applied these montages on the 25 individual heads to compare the focality and intensity they achieved to those achieved by the individually optimized montages. Note for 2 of these 6 targets, no montage was found in the literature. The montages we found in the literature are summarized in Table 1.

**Table 1:** Montages of 2-pair TI found in the literature targeting the brain regions studied in this paper. 1 mA current was applied at each pair of electrodes.

| Target location | Right hippocampus | Left DLPFC | Left motor cortex | Right amygdala | Right caudate | Left thalamus |
| --- | --- | --- | --- | --- | --- | --- |
| Montage and reference | Fp1–FT8, P7–TP8;<br>Inferred from Violante et al., 2023 | N/A | F1–CP1, FC1–CP3 (von Conta et al., 2021) | N/A | F3–F4 and TP7–TP8 (Vassiliadis et al., 2024a) | F7–PO7, F8–PO8 (von Conta et al., 2021) |

### Sensitivity analysis on the optimized montages

To find out how sensitive each stimulation modality is with respect to the optimized montages, we ran a simulation study on Subject 1. We randomly generated 1000 montages for each of the three stimulation modalities at each target. The random montage was generated by “randperm” function in Matlab, and had the same number of electrodes as the optimized montage (4 for 2-pair TI, and K for 2-array TI and HD-TES, where K is the number of electrodes in the optimized montage that have a dosage higher than 10^-6^ mA). The current at each electrode in the random montage was generated also randomly by the “rand” function in Matlab. The sum and L1-norm of the random montage was made to be 0 and 4 mA, respectively, to be consistent with the optimized montage. We applied these 1000 random montages along with the optimized montages to the model and compared the generated modulation depth (MD) and its focality at the target. For the optimized montages of HD-TES and 2-array TI, we used the strictest energy constraint at which the injected current just reaches the total budget of 2 mA, i.e., the left-most point on the focality-modulation curve. To compare the focality, we computed the loss of focality from using random montages compared to using the optimized montages, where positive value means focality loss and negative means focality gain. For TI, we also simulated a linear version of TI where the non-linear relation between the induced MD and the lead field was replaced by a simple sum (Huang and Parra, 2019), i.e., MD**(r)** = 2min(|**d(r)**^T^**A(r)s_1_**|,|**d(r)**^T^**A(r)s_2_**|) was replaced by MD**(r)** = |**d(r)**^T^**A(r)s_1_** + **d(r)**^T^**A(r)s_2_**|, where **A(r)** is the lead field at brain location **r**, **d(r)** is the unit directional vector at **r**, and **s_1_** and **s_2_** are the montage vectors for the two frequencies in the optimized TI montages. To rule out the effect of the specific head anatomy of this subject, we repeated this sensitivity analysis on a different model of Subject 1 that has a homogeneous conductivity. To make this model, we ran ROAST on Subject 1 for the lead field and specified all the six tissues to have the same conductivity of 1.0 S/m (gel and electrode were kept the default conductivities). Then we ran the same optimization algorithms for HD-TES, 2-array and 2-pair TI, to maximize the stimulation focality at the 6 targets, following the same procedure as described above. These optimized montages were compared to 1000 random montages at each target following the same procedure.

### Phantom validation of the individually optimized montages

To test if our optimized 2-pair TI indeed outperforms the montages found in the literature and random montages, we created a head phantom using the MRI of Subject 1 and recorded the stimulation strength at the target during 2-pair TI. Specifically, the stereolithography (STL) file of the outer scalp surface of Subject 1 generated by ROAST was used to 3D print a mold (Fig. 2A). 3D coordinates were obtained from the MRI to locate a tube in the radial-in direction pointing from the scalp surface to the location of the right hippocampus (Target 1). The tube was 3D printed together with the head mold (Fig. 2B). About 375 grams of agar powder and 2750 ml saline were mixed and heated to boiling temperature on a stove. The mixture was stirred during the heating process to prevent agglomeration. Once evenly mixed, the agar solution was poured into the mold to fill it up. The entire assembly was air-cooled at room temperature for 1 hour and was then placed in a refrigerator overnight to allow complete cooling. The mold pieces were then separated and removed to harvest the agar phantom (Fig. 2C). A strip of 8-contact stereoelectroencephalogram (sEEG) electrodes (Ad-Tech, Oak Creek, WI) was then inserted into the tube (Fig. 2D) until the tip of the strip reached the end of the tube where the right hippocampus is. The remaining empty space of the tube was then filled up by the same agar solution using a syringe. The sEEG electrode strip was attached to PowerLab data acquisition platform and the signal was recorded by LabChart software (ADInstruments, Colorado Springs, CO) running on a local workstation. An EEG cap with 10/10 electrode layout was placed on the phantom to mark the electrode positions that will be used for stimulation. The cap was then removed, and stimulation electrodes were placed on the phantom by sticky tapes. The stimulation electrodes were connected to our HD-IFS device (Soterix, Woodbridge, NJ) to perform the 2-pair TI stimulation on the head phantom. The whole setup is shown in Fig. 2E for one montage.

**Fig. 2:**
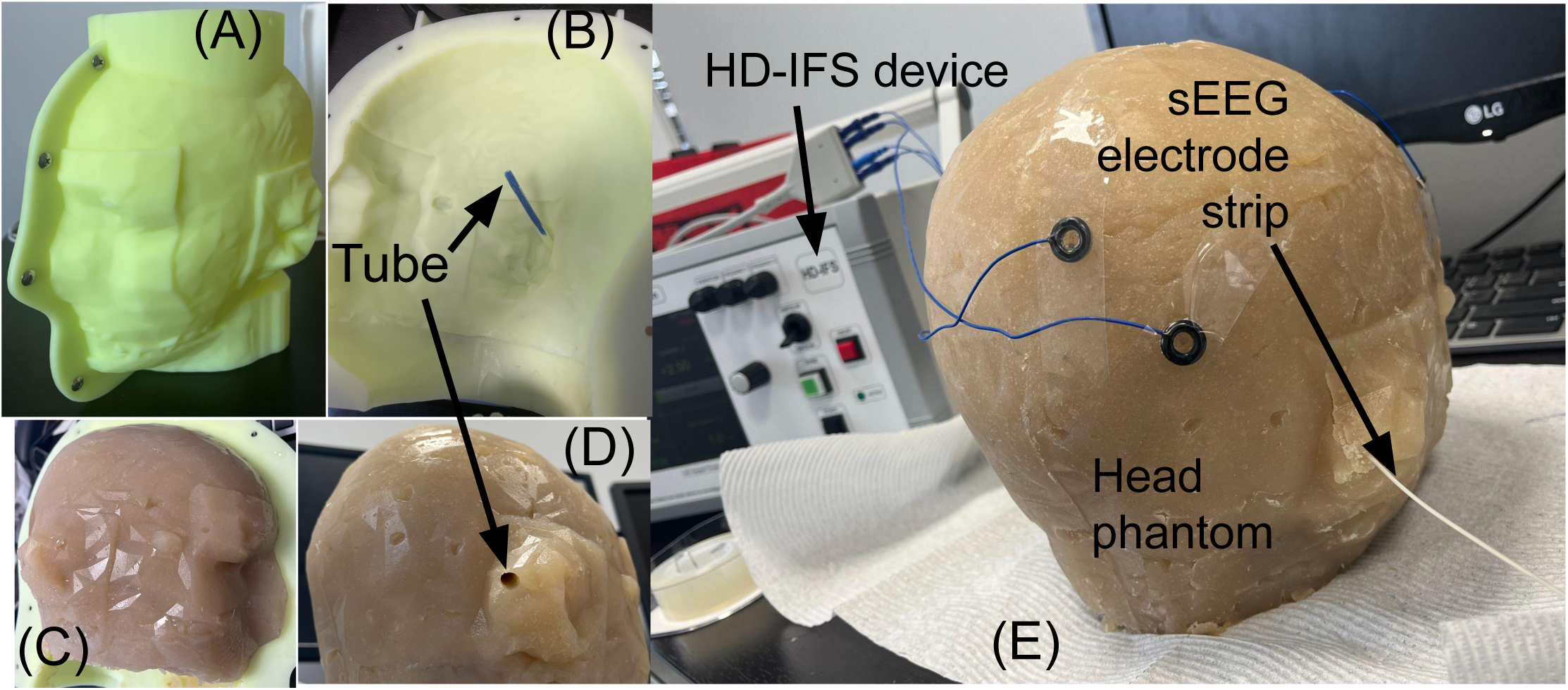
(A) 3D printed mold replicating the geometry of the scalp of Subject 1. (B) Mold seen from inside highlighting a tube printed with the mold indicating the radial direction from scalp surface to the right hippocampus. (C) Right lateral view of harvested head phantom made of agar powder, with the tube indicated by the arrow in (D). The whole experiment setup is shown in (E).

Before starting off the recording sessions described below, bad electrode contacts were checked and fixed. We first set the frequency for both channels to be the same to confirm the system is quasistatic under different stimulation intensities (0.1 – 5 mA) and frequencies (1 – 2,000 Hz). We recorded under the optimized montage that achieved the best focality in the model at the right hippocampus on Subject 1. As sEEG electrodes can only record data at 8 locations in the head, it cannot collect enough spatial data to compute the 3D focality as in the model. The advantage of the max-focality montage may not be captured by our recording setup. Therefore, in the model, we also searched for the montage that maximized the MD in the radial-in direction at the target, regardless of focality, as the radial MD is the only quantity that can be measured by our setup. Note we performed this optimization on the model of Subject 1 with the homogeneous conductivity (see section on sensitivity analysis) as the head phantom is made up of only one material. We also recorded the data under montages found in the literature (Violante et al., 2023) and random montages. Table 2 summarizes the montages we collected data for. For each of the columns in Table 2, we also altered the current applied at each pair of electrodes (number in the parentheses) while keeping the electrodes the same. This is to test if the optimized montages are sensitive to small changes, and the 1:1 vs 1:3 current ratios reported in Violante et al., 2023.

**Table 2:** Montages of 2-pair TI targeting the right hippocampus tested in the phantom experiment. Number in parentheses indicates the dosage of current (in mA) applied at that pair of electrodes.

| Optimal focality montage | Optimal MD montage | Literature montage (Inferred from Violante et al., 2023) | Random montage |
| --- | --- | --- | --- |
| Fp1–AF8 (1.35), PO4–PO8 (0.65) | T8–F1 (1.05), TP8–C3 (0.95) | Fp1–FT8 (1), P7–TP8 (1) | AF3–C3 (1), C4–PO4 (1) |
| Fp1–AF8 (0.2), PO4–PO8 (1.8) | T8–F1 (0.2), TP8–C3 (1.8) | Fp1–FT8 (0.5), P7–TP8 (1.5) | AF3–C3 (0.5), C4–PO4 (1.5) |

For all the recording sessions, we used bipolar sinusoidal waveform, with 15 s ramp up, 15 s ramp down, 1000 Hz and 1010 Hz as the two interfering frequencies. Each session lasted 1 min excluding the time ramping up and down. Sampling rate was set as 4000 Hz. Raw data was saved in Matlab format and was first notch filtered to remove the 60-Hz power interference. The first and last second of data in each session were discarded to ensure data quality. Electric field at the sEEG electrodes was then calculated by subtracting voltages along adjacent sEEG electrodes and dividing by the electrode distance (5 mm). The envelope of the electric field was then computed by applying the Hilbert transform (Huang and Parra, 2019). The raw envelope contains noisy frequencies, and thus was bandpass filtered at 10 Hz beat frequency (difference between the two frequencies used). MD was then calculated as the difference between the peak and valley of the envelope signal. MDs at the sEEG contacts were plotted and compared across the montages we tested. Recorded MDs were also compared against the predicted MDs from the homogeneous-conductivity model of Subject 1. To read out modeled values, a line going in radial-out direction from the right hippocampus to the cortical surface was generated. Coordinates along this line were used to extract the MD from the model.

## Results

### Inter-individual variability of optimized TI and HD-TES

Fig. 3 shows the optimized montages for targeting the right hippocampus in the first five subjects, at the smallest energy constraint where the injected current just reaches 2mA. The left column (A1-E1), middle column (A2-E2), and right column (A3-E3) shows the optimized montages from conventional HD-TES, 2-array TI, and 2-pair TI, respectively. For these five subjects, the injected current as a function of electrode index (from #1 to #72) is also shown in panels F1–F3. It is evident that the optimal montages vary significantly across different subjects for targeting the same location in the brain, for all three stimulation modalities, due to inter-individual variabilities in head anatomy and tissue conductivity. We note some interesting patterns on the spatial distribution of stimulation electrodes for the 2-array TI: one frequency tend to concentrate on the right scalp (panels A2–E2, right topoplots), and for the 2-pair TI, the two frequencies are separated at the left and right scalps, with the locations of the electrodes on the left scalp changing significantly across subjects (panels A3–E3). We found similar patterns on the other 20 subjects.

**Fig. 3:**
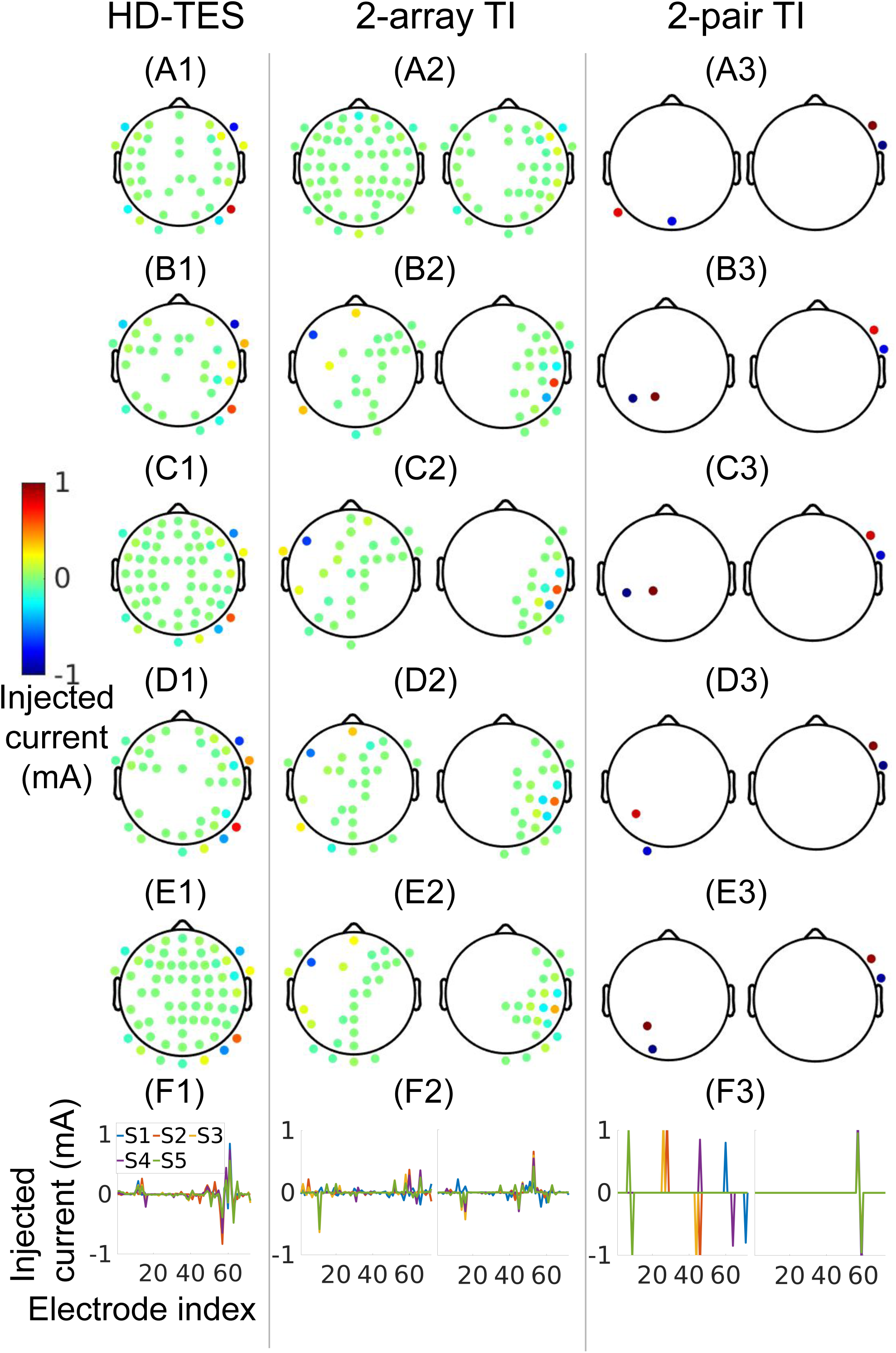
Topoplots of optimized montages targeting the right hippocampus for HD-TES (left column), 2-array TI (middle column), and 2-pair TI (right column). Here only the montages for the first five subjects are shown (rows A–E, respectively). For TI, the two topoplots at each panel show the montages for the two frequencies. As the injected current at most electrodes is close to zero for HD-TES and 2-array TI, it is also shown as a function of the electrode index (1– 72) in row F for these five subjects (S1–S5).

The standard deviation of these injected current across the 25 subjects at each electrode is shown in Fig. 4, for the 6 target locations: right hippocampus (row A1–A3), left DLPFC (row B1– B3), left motor cortex (row C1–C3), right amygdala (row D1–D3), right caudate (row E1–E3), left thalamus (row F1–F3), and the three stimulation modalities: conventional HD-TES (column A1-F1), 2-array TI (column A2-F2), 2-pair TI (column A3-F3). Note that the maximal standard deviation is 0.90 mA (panel F3).

**Fig. 4:**
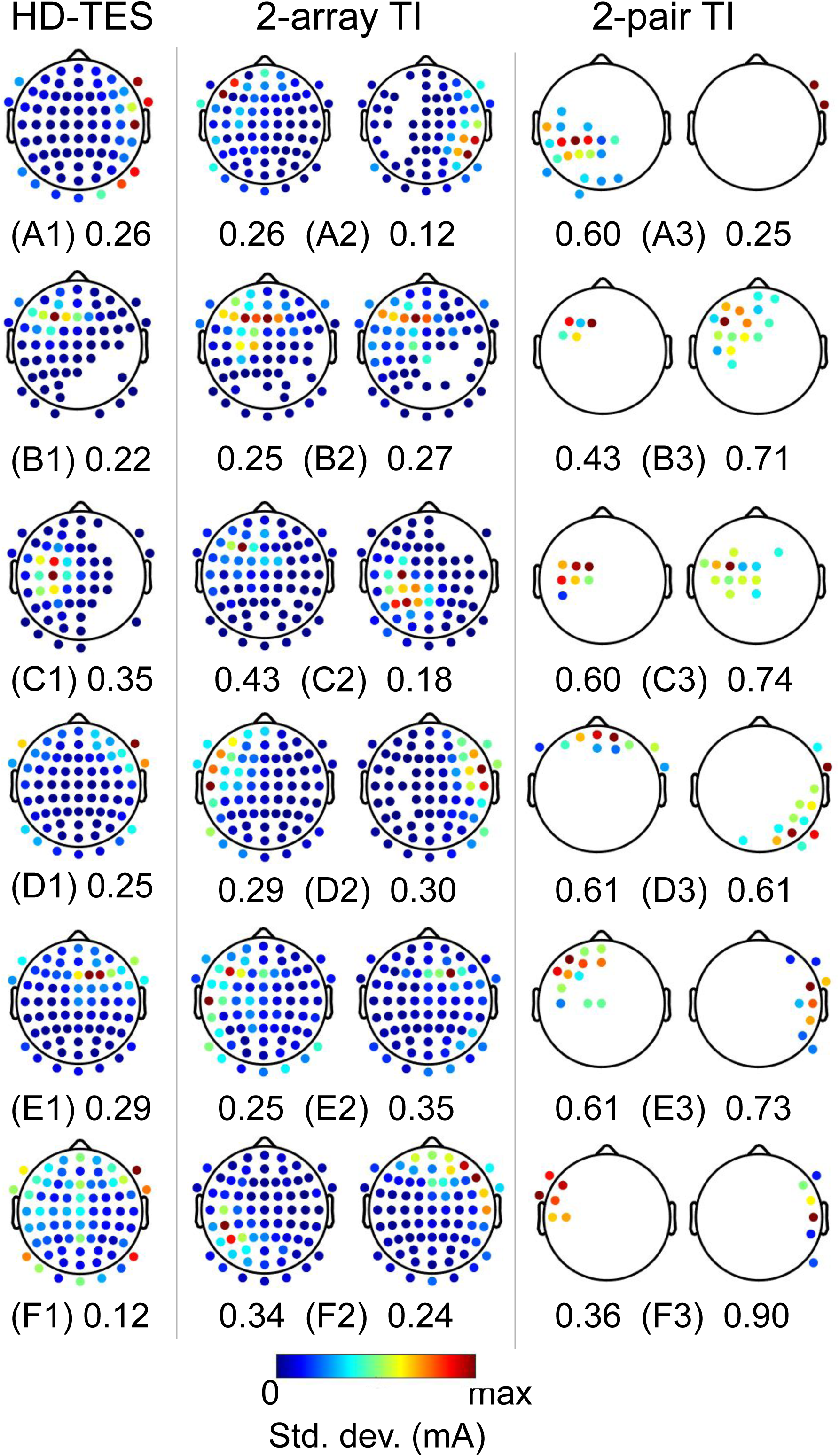
Topoplots of standard deviation across the 25 subjects of the injected current at each electrode, for HD-TES (left column), 2-array TI (middle column), and 2-pair TI (right column), in targeting the right hippocampus (row A), the left DLPFC (row B), the left motor cortex (row C), the right amygdala (row D), the right caudate (row E), and the left thalamus (row F). For TI, the two topoplots at each panel show the standard deviation for the two frequencies. Colorbar represents the standard deviation and runs from 0 to the maximal value of each topoplot. The corresponding maximal standard deviation for each plot is noted beside the panel label.

The inter-individual variability can also be seen in the focality-modulation curves shown in Fig. 5. Different subjects have the curves residing in different locations in the focality-modulation plane. For the sake of comparison across subjects, we normalized the modulation depth into the range of 0% and 100% by linear interpolation. We see that under the same level of modulation depth, the focality at the target varies across subjects. For example, at the normalized modulation depth of 80%, focality of HD-TES has a standard deviation of 0.40 cm for Target 1 (Fig. 5B1, blue shaded curve), 0.49 cm for Target 2 (Fig. 5B2), 0.87 cm for Target 3 (Fig. 5B3), 0.32 cm for Target 4 (Fig. 5B4), 0.89 cm for Target 5 (Fig. 5B5), and 0.36 cm for Target 6 (Fig. 5B6). For 2-array TI, this standard deviation is 0.65 cm (Fig. 5B1, red shaded curve), 0.54 cm, 0.83 cm, 0.37 cm, 1.20 cm, and 0.57 cm for the 6 targets, respectively. For 2-pair TI, as one cannot explicitly specify any energy constraint in the optimization algorithm, one therefore obtains different modulation depths on different subjects, and focality was compared at these varying modulation depths. The standard deviation of the optimized focality is 0.26 cm (Fig. 9A1, circles), 0.59 cm (Fig. 9A2), 0.93 cm (Fig. 9A3), 0.35 cm (Fig. 9A4), 0.84 cm (Fig. 9A5), and 0.60 cm (Fig. 9A6) for the 6 targets, respectively.

**Fig. 5:**
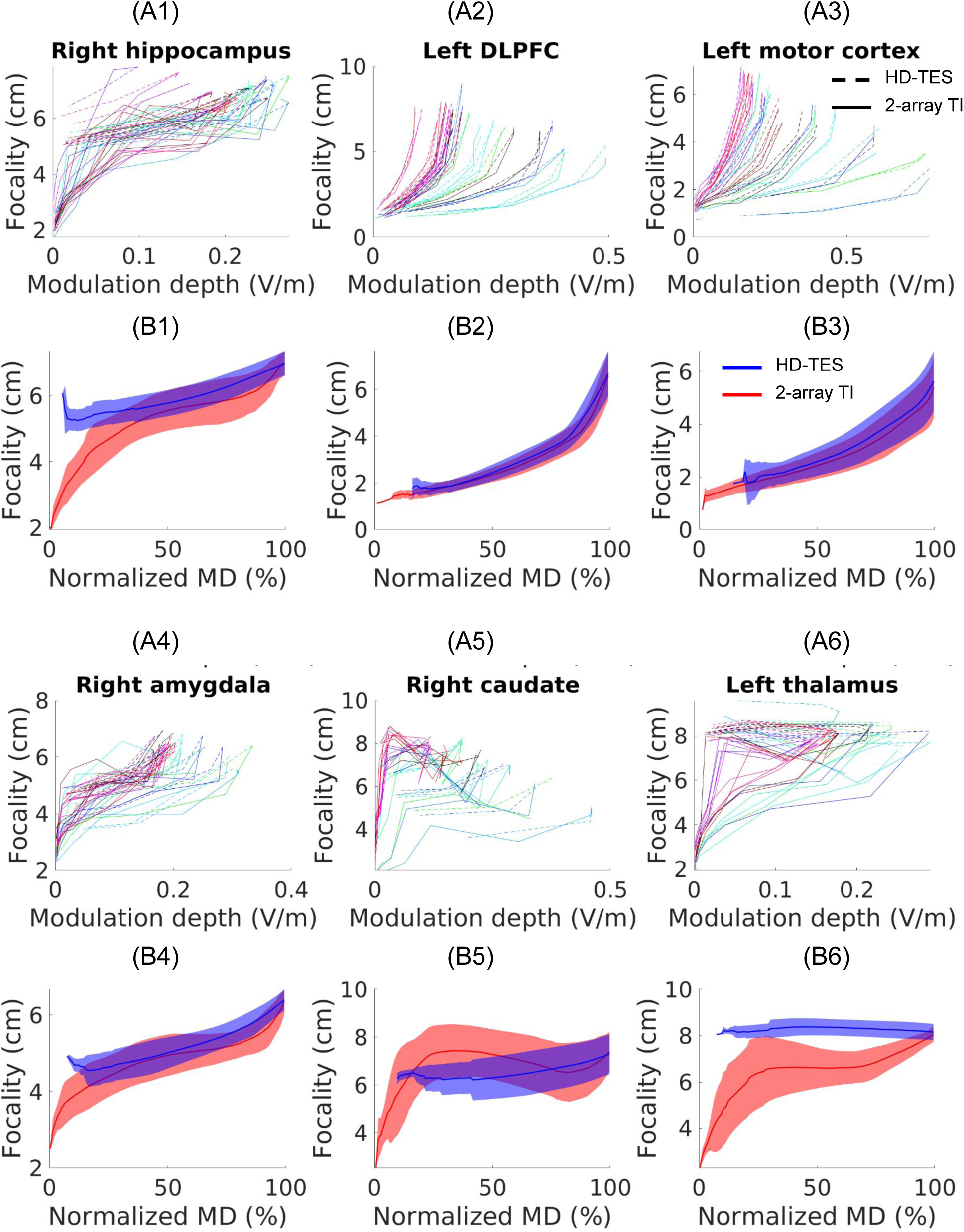
Focality-modulation curves for the 25 subjects targeting the right hippocampus (A1), the left DLPFC (A2), the left motor cortex (A3), the right amygdala (A4), the right caudate (A5), and the left thalamus (A6), under 12 different levels of energy constraints for the non-target area. Different colors represent different subjects, with solid lines representing 2-array TI and dashed lines representing HD-TES. Normalized curves are shown in panels B1–B6, where modulation depth (MD) was normalized between 0% and 100%. Solid lines and shaded areas represent the mean and standard deviation of the focality measures (in centimeter) across subjects, with red and blue indicating 2-array TI and HD-TES, respectively.

### Optimized 2-array TI achieves higher focality than optimized HD-TES in all individuals at most targets

At the same level of modulation depth, optimized 2-array TI and optimized HD-TES achieve different focalities, for all the 25 individual heads (Fig. 5). A two-way (modulation, method) repeated measures ANOVA shows that this difference between TI and HD-TES is significant (Fig. 5, B1–B6; for the 6 targets: F(1,63)=27.61, p<10^-4^; F(1,61)=56.30, p<10^-7^; F(1,54)=77.60, p<10^-8^; F(1,71)=13.09, p=0.001; F(1,58)=18.31, p<10^-3^; F(1,60)=152.70, p<10^-11^). At the 80% modulation, a pair-wise t-test shows that TI has significantly better focality than HD-TES for the 6 targets except Target 5: t(24)=4.98, p<10^-4^; t(24)=3.24, p=0.004; t(24)=6.20, p<10^-5^; t(24)=8.19, p<10^-7^; t(24)=1.43, p=0.17; t(24)=12.14, p<10^-11^. For Target 5 (Fig. 5B5), advantage of TI over HD-TES in focality is only slightly significant at the 90% modulation level (t(24)=2.11, p=0.046); at a lower modulation level of 30%, TI actually achieves worse focality than that from HD-TES at Target 5 (t(21)=-11.37, p<10^-9^). Therefore, different from our previous finding on the MNI152 head (Huang et al., 2020), optimized 2-array TI may not always perform better than optimized HD-TES in terms of stimulation focality. This may be due to the specific location of Target 5, and the numerical instability and sensitivity to initialization of the optimization algorithm at stricter energy constraints we noticed before (Huang et al., 2020). Nevertheless, 2-pair TI still achieves better focality at Target 5 (in average 4.64 cm) compared to HD-TES (higher than 5 cm), see Fig. 9A5 (circles) and Fig. 5B5 (blue shaded curve).

To visualize this difference, Fig. 6 shows some examples. Fig. 6A is from Subject 3 at the 4th energy constraint for targeting the right hippocampus, Fig. 6B is from Subject 3 at the 8th energy constraint for targeting the left DLPFC, and Fig. 6C is from Subject 6 at the 9th energy constraint for targeting the left motor cortex. Here subjects and energy constraints were chosen arbitrarily to just show the examples. We slightly adjusted the energy constraint for HD-TES so that it generates the same modulation at the target as that from 2-array TI. The boost in focality in TI is more evident in the deep target than the cortical target (e.g., 2.05 cm boost in the right hippocampus (Fig. 6, panels A2&A6) vs. 0.19 cm improvement in the left motor cortex (Fig. 6, panels C2&C6), which is consistent with our previous finding on the MNI152 head (Huang et al., 2020). Note that more electrodes are needed for targeting deeper structures (Fig. 6, A1, A4, A5 vs. C1, C4, C5).

**Fig. 6:**
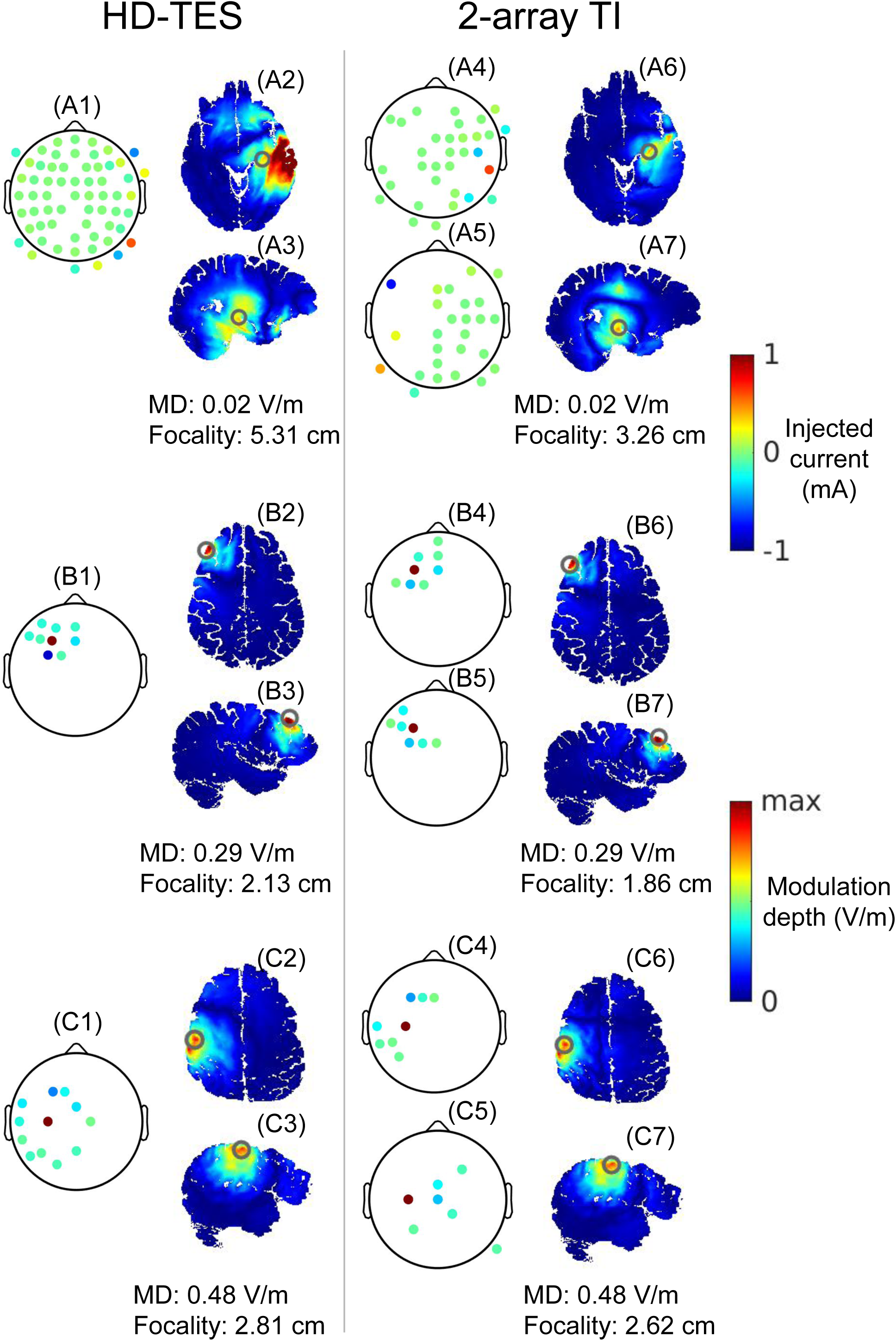
Examples of optimized montages and the corresponding distributions of modulation depth for targeting the right hippocampus (A1–A7), the left DLPFC (B1--B7), and the left motor cortex (C1–C7). Panels to the left of the vertical divider show results from optimized HD-TES, while panels to the right of the divider show optimized 2-array TI. For TI, the two topoplots show the montages for the two frequencies. Modulation depth (MD) and the focality of the MD at the target are noted.

### Individually optimized 2-array TI montages give higher focality than that from common montages at most targets

Fig. 7 shows the focality-modulation curves for all the 25 subjects from using individually optimized montages and a common montage obtained from the standard MNI152 head targeting the 6 locations being studied, using conventional HD-TES. Interestingly, individually optimized and common montages do *not* give significantly different results (Fig. 7, B1–B6; two-way repeated measures ANOVA for the 6 targets: F(1,63)=1.81, p=0.19; F(1,61)=0.26, p=0.61; F(1,55)=1.14, p=0.30; F(1,71)=3.76, p=0.06; F(1,57)=0.29, p=0.60; F(1,60)=0.01, p=0.94).

**Fig. 7:**
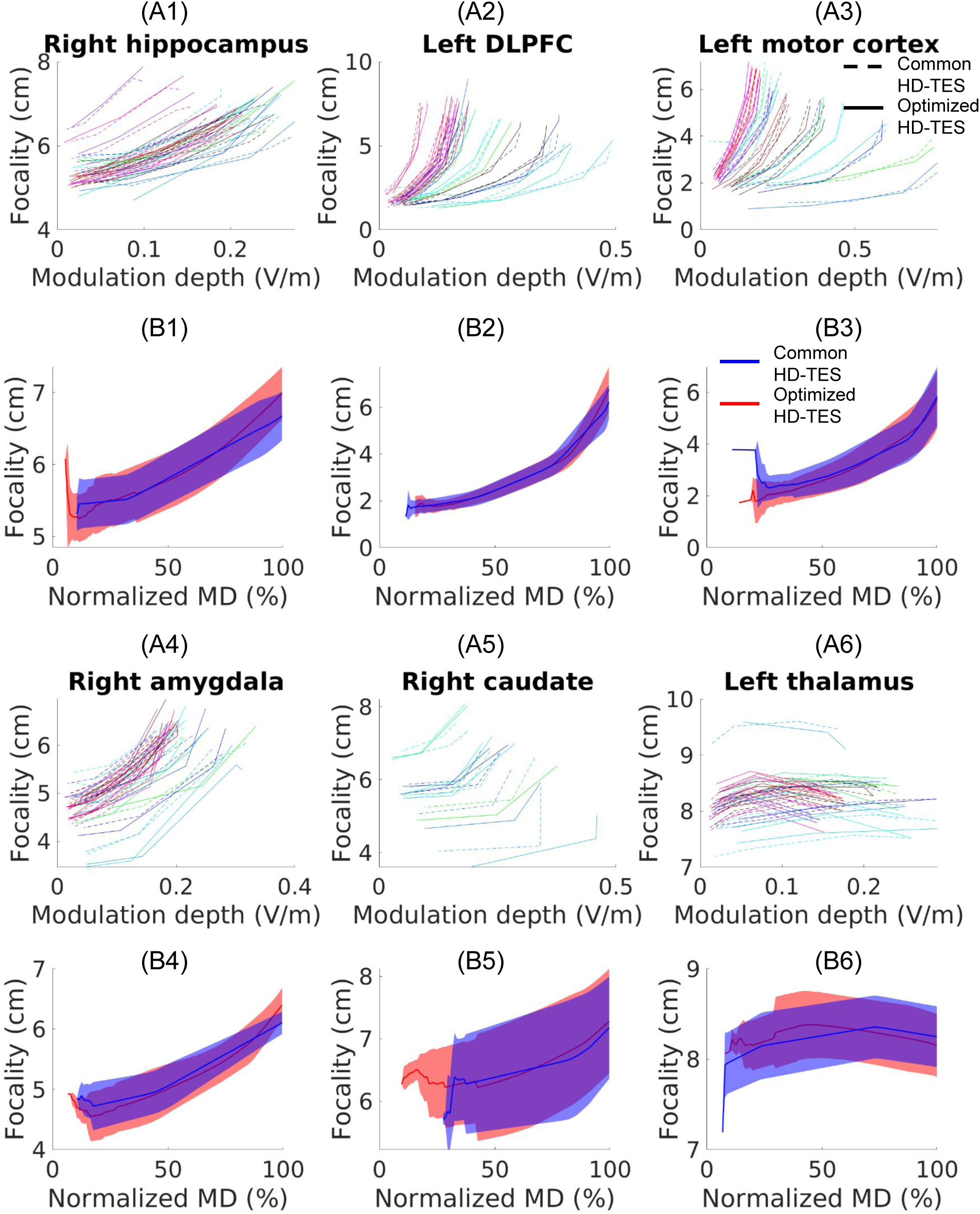
Focality-modulation curves for the 25 subjects targeting using HD-TES the right hippocampus (A1), the left DLPFC (A2), the left motor cortex (A3), the right amygdala (A4), the right caudate (A5), and the left thalamus (A6), under 12 different levels of energy constraints for the non-target area. Different colors represent different subjects, with solid lines representing results using individually optimized montages and dashed lines results using a common montage obtained from targeting on the standard MNI152 head. Normalized curves are shown in panels B1–B6, where modulation depth (MD) was normalized between 0% and 100%. Solid lines and shaded areas represent the mean and standard deviation of the focality measures (in centimeter) across subjects, with red and blue indicating optimized and common montages, respectively.

However, this is not true for the 2-array TI, where individually optimized montages give significantly different focalities compared to those from using a common montage, for all the targets except Target 4 (Fig. 8, B1–B6; two-way repeated measures ANOVA for the 6 targets: F(1,83)=42.40, p<10^-6^; F(1,63)=14.97, p<10^-3^; F(1,57)=71.18, p<10^-7^; F(1,89)=0.01, p=0.93; F(1,83)=12.98, p=0.001; F(1,71)=26.16, p<10^-4^). The non-siginificance found on Target 4 may be due to the limited availability of electrodes placed at the lower part of the scalp (see Discussions) that limited the full beamforming potential of optimized 2-array TI, as Target 4 is the deepest among the 6 targets. At the normalized modulation depth of 80%, compared to using the individually optimized montages, using a common montage drops the focality in average 0.77 cm for Target 1 (t(24)=6.79, p<10^-6^), 0.29 cm for Target 2 (t(24)=3.28, p=0.003), and 0.35 cm for Target 3 (t(24)=5.24, p<10^-4^). For Targets 5 and 6 (Fig. 8, B5&B6), using a common montage does not seem to affect the focality at the 80% modulation (t(24)=-0.23, p=0.82; t(24)=2.10, p=0.05), but it does significantly deteriorate the focality at 50% modulation (average drop of 0.71 cm for Target 5 (t(24)=5.92, p<10^-5^) and 1.05 cm for Target 6 (t(24)=6.09, p<10^-5^)).

**Fig. 8:**
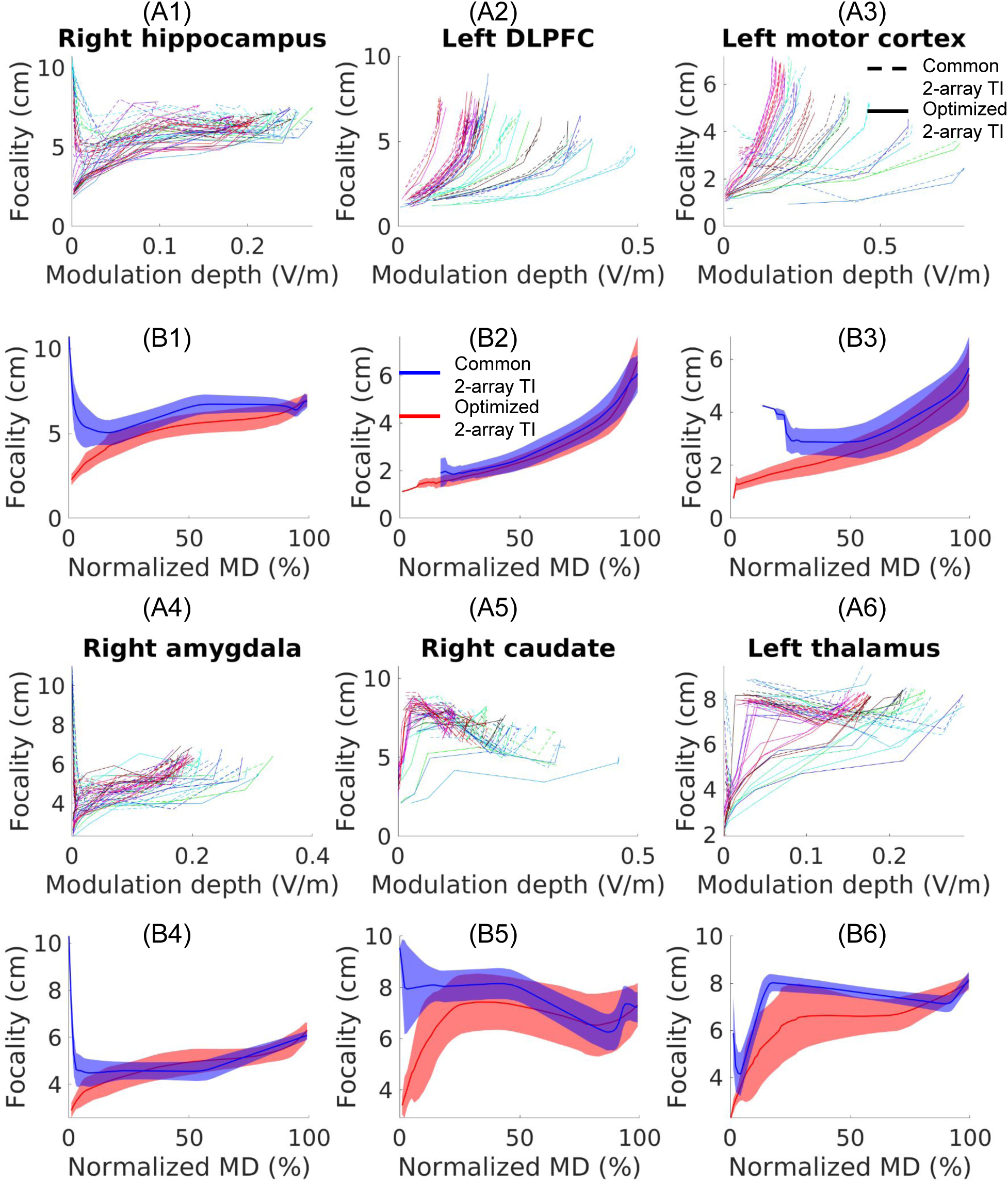
Similar to Fig. 7, but for 2-array TI.

**Fig. 9:**
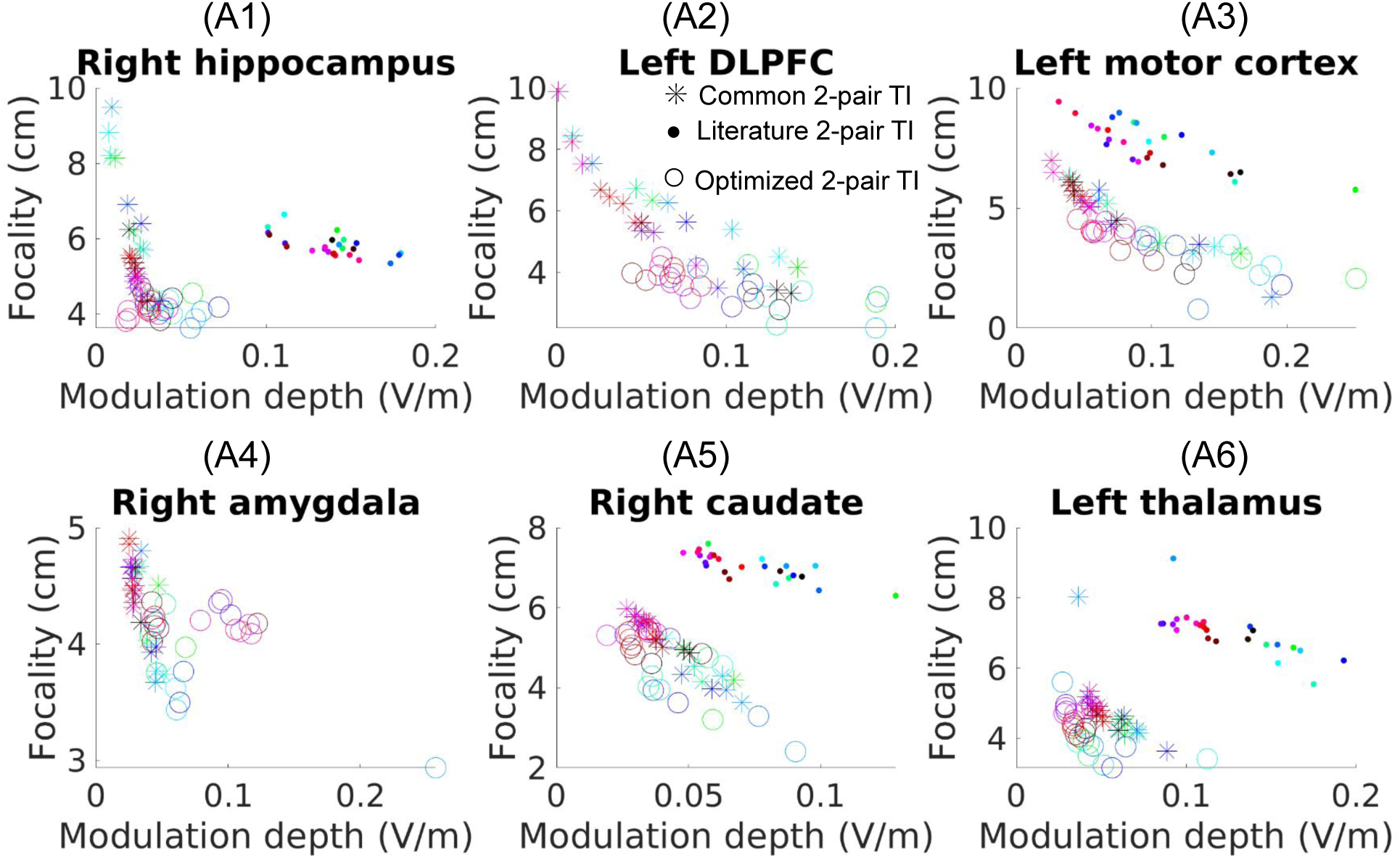
Similar to Fig. 7, but for 2-pair TI. Note there is no energy constraint explicitly specified in optimizing TI using two pairs of electrodes, and thus the results are shown in points instead of curves. Different colors represent different subjects. Results from individually optimized montages and optimized montages from the standard MNI152 head are represented by circles and asteroids, respectively. We also show the results obtained from montages found in the literature targeting these brain regions (if available), indicated by dots.

### Individually optimized 2-pair TI montages give higher focality than that from common montages and literature montages

For the more popular 2-pair TI, we observe an even more obvious difference between using individually optimized montages and a common montage (Fig. 9, circles vs. asteroids). Using a common montage on individual heads without optimization leads to a significantly worse focality at the target (focality drops in average: 1.69 cm at Target 1 (p<10^-5^); 2.33 cm at Target 2 (p<10^-7^); 1.65 cm at Target 3 (p<10^-9^); 0.38 cm at Target 4 (p<10^-9^); 0.37 cm at Target 5 (p<10^-6^); 0.49 cm at Target 6 (p<10^-4^)). The common montage also generates weaker modulation strength for the first 4 targets (MD drops in average: 0.02 V/m at Target 1 (p<10^-4^); 0.04 V/m at Target 2 (p<10^-4^); 0.04 V/m at Target 3 (p<10^-5^); 0.05 V/m at Target 4 (p<10^-4^)). The common montage however induces a slightly higher (0.01 V/m) modulation at Target 6 (p<10^-4^), and does not significantly affect the modulation at Target 5 (p=0.28). Literature montages (Table 1) also significantly drop the focality compared to the individually optimized ones (Fig. 9, dots vs. circles; focality drop in average: 1.66 cm at Target 1 (p<10^-21^); 4.41 cm at Target 3 (p<10^-15^); 2.40 cm at Target 5 (p<10^-14^); 2.74 cm at Target 6 (p<10^-22^)), even though the literature montages induce higher stimulation at most targets (MD increase: 0.10 V/m at Target 1 (p<10^-17^); 0.03 V/m at Target 5 (p<10^-8^); 0.08 V/m at Target 6 (p<10^-14^)). All p values are from pair-wise t-test.

As some examples, Fig. 10 visualizes the difference between using optimized and common montages in targeting the right hippocampus, for the three stimulation modalities (HD-TES, 2-array TI, and 2-pair TI). Fig. 10A is Subject 3 at the 5th energy constraint, and Fig. 10B is Subject 3 at the 4th energy constraint. Again subjects and energy constraints were chosen arbitrarily to just show the examples. Fig. 10C is Subject 3 under TI using two pairs of electrodes. Note using common montage for TI did not output the same modulation as it was obtained from the standard MNI152 head instead of the head being studied. We noticed a small drop in focality when using a common montage for HD-TES (0.34 cm, Fig. 10, panels A5&A2), but a 10 times big drop in 2-array TI (3.40 cm, Fig. 10, panels B7&B3), and also an almost 7 times drop in 2-pair TI (2.29 cm, Fig. 10, panels C7&C3).

**Fig. 10:**
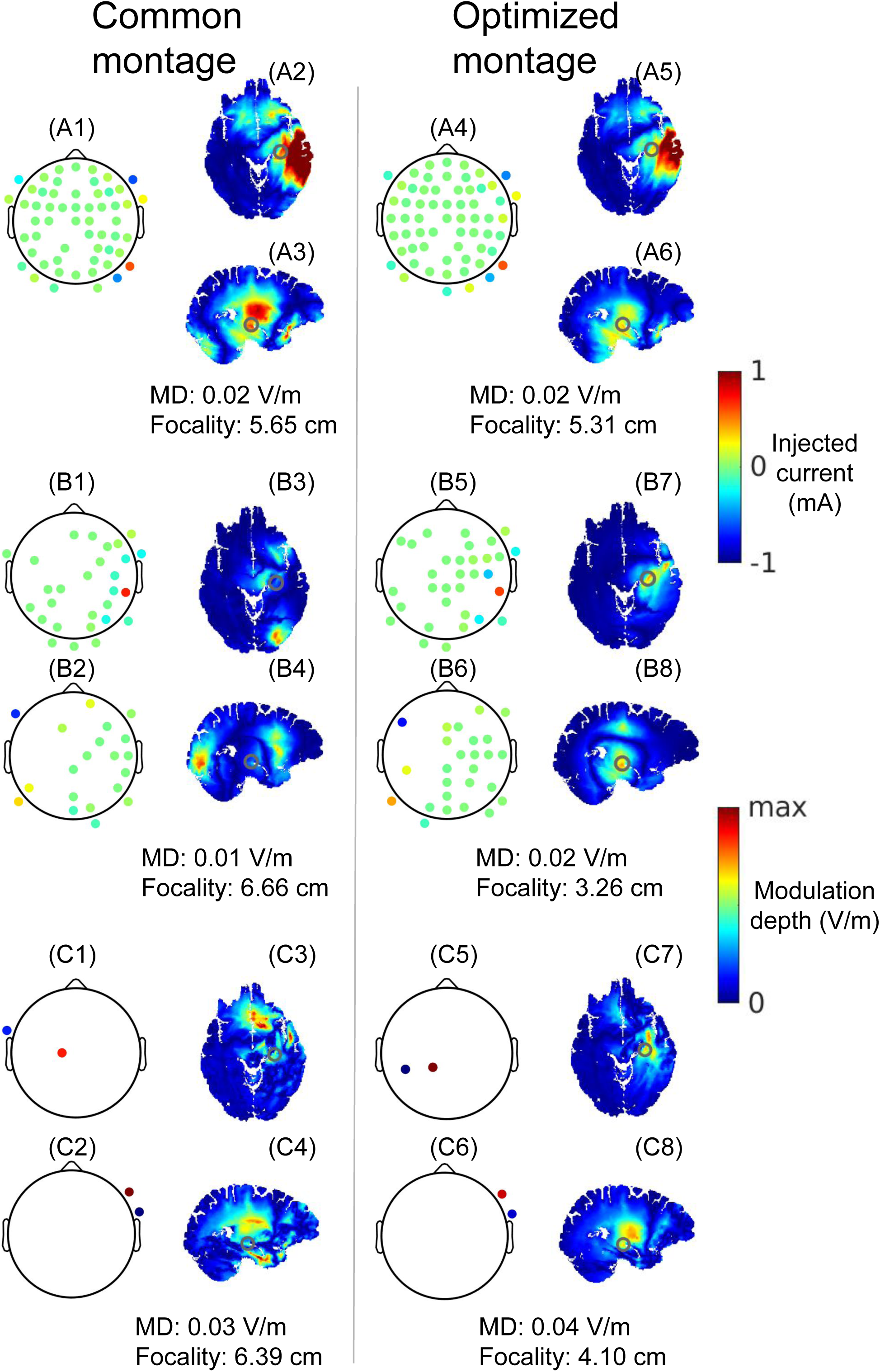
Examples of montages and the corresponding distributions of modulation depth for HD-TES (A1–A6), 2-array TI (B1--B8), and 2-pair TI (C1–C8). Panels to the left of the vertical divider show results from using a common montage obtained from targeting the right hippocampus on the standard MNI152 head, while panels to the right of the divider show results from using individually optimized montages targeting the right hippocampus. For TI, the two topoplots show the montages for the two frequencies. Modulation depth (MD) and the focality of the MD at the target are noted.

### Sensitivity analysis on the optimized montages

Fig. 11 shows the results of the sensitivity analysis performed on Subject 1. Fig. 11A is the focality-modulation plot for all the montages used for Target 1 (right hippocampus). Optimized montages are represented in red symbols, while random montages are in blue (for readability only results from 10 random montages are shown). It is clear that one loses significant focality when random montages are used, for all the three stimulation modalities: 2-pair TI (asteroids), 2-array TI (plus signs), and HD-TES (crosses). Focality is not sensitive to montages any more if the non-linearity of TI physics is removed (Fig. 11A, circles and triangles). Fig. 11B shows the sensitivity of focality (positive value: loss; negative: gain) at all the targets using the 1000 random montages. We see that one can lose focality up to 9.27 cm when using random montages (2-array TI on Target 3), but this sensitivity of focality to montage changes drops significantly when the non-linearity is removed (2-array TI: yellow vs magenta violins; 2-pair TI: blue vs red), though the drop is small at Target 6 (left thalamus) for 2-array TI. This suggests that the sensitivity of focality to the TI montages may come from the non-linear relation between the MD and the lead field (Huang and Parra, 2019). It is interesting to note that 2-pair TI is not always more sensitive than HD-TES: at shallow targets (left DLPFC & left motor cortex), HD-TES appears more sensitive than 2-pair TI (green vs blue violins), while in the deep locations (the other 4 targets), 2-pair TI is more sensitive than HD-TES. Sensitivity of HD-TES and 2-array TI are on par and do not show this depth-dependent pattern (green vs yellow). We also found that TI sensitivity drops significantly when TI non-linearity is removed on the model of this same subject but with homogenous conductivity values (figure not shown), ruling out the possibility that a specific tissue layer contributes to the TI sensitivity. Note Fig. 11 shows the differences in focality between optimized and 1000 random montages, while Figs. 7-9 are differences between optimized and one common montage *optimized* from the standard MNI152 head. So we observe a bigger difference in Fig. 11. These data suggest that focality of stimulation would drop significantly if a random montage is adopted. If an individually optimized montage is not available, one at least should use an optimized montage obtained from the MNI152 head.

**Fig. 11:**
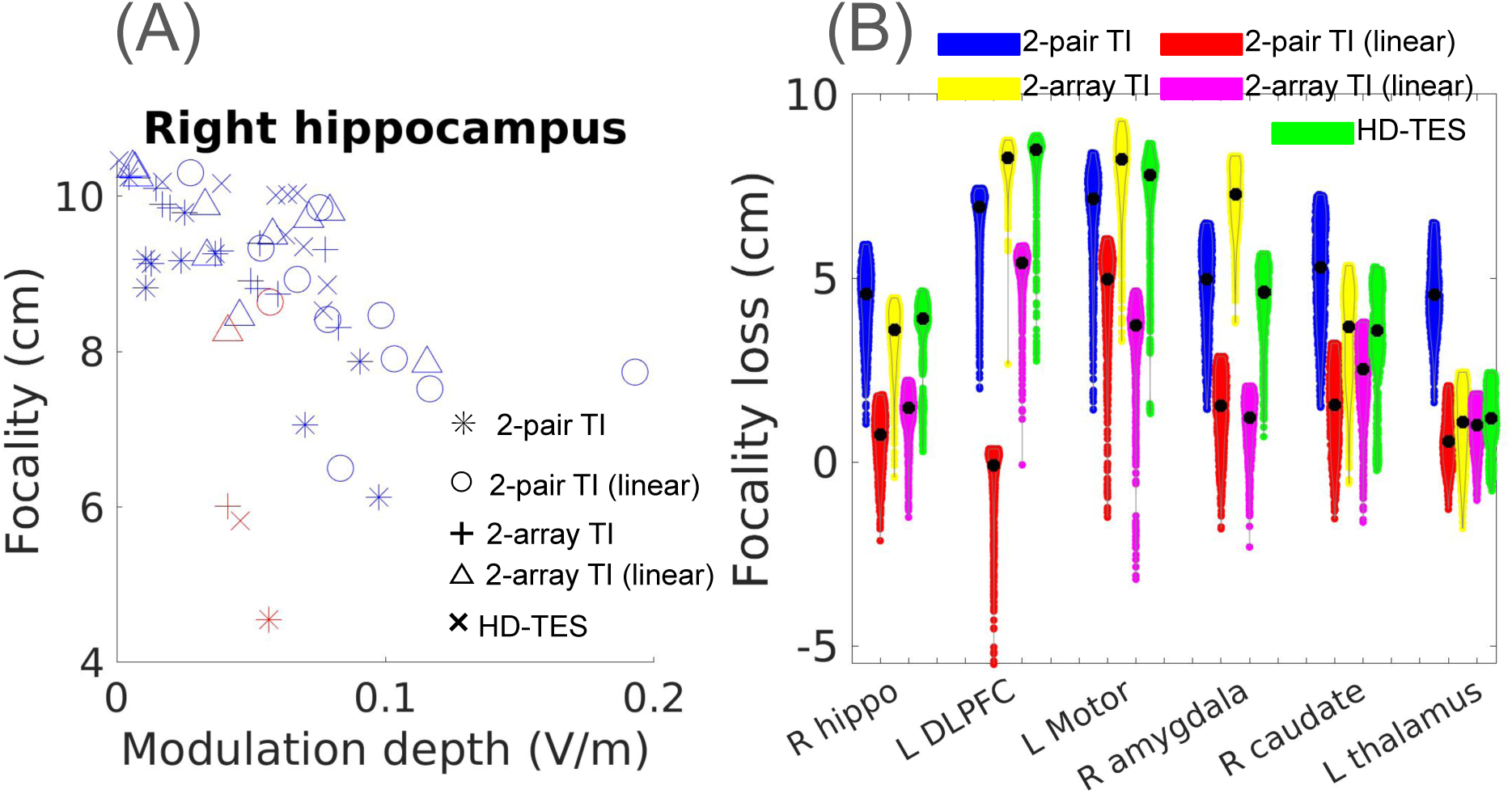
(A) Focality-modulation plot for Subject 1 from the sensitivity analysis. Results from optimized montages are in red symbols, and results from random montages are in blue (only 10 out of 1000 random montages are randomly selected to be plotted here for better readability). The linear version of TI (circles and triangles) adopts the same montage as TI but the non-linearity is replaced by a linear sum (see Methods for details). (B) Violin plots of focality loss (i.e., sensitivity of focality) when 1000 random montages are used on Subject 1 instead of optimized montages, for all the targets and stimulation modalities studied. Black dots in violin are the median values.

### Head phantom validation of optimized montages

Fig. 12A shows the MD recorded at the sEEG contacts placed around the right hippocampus in the head phantom of Subject 1. Contact #1 at the tip of the sEEG strip was placed at the MNI coordinates of the right hippocampus. It is clear that the montages optimized for focality (optFocality & optFocality2) induced very weak MD, even weaker than that from the literature montages, and close to the MD from random monages. The strongest MD is from the montage that specifically maximized the MD at the target (optMD), and changes in dosage in this optimized montage greatly reduced the MD (optMD2), consistent with the *in silico* sensitivity analysis. Literature montages (literature & literature2) from Violante et al., 2023 induces weaker MD than the optimized montage (optMD). 1:3 current ratio (literature2) gives weaker MD than 1:1 current ratio (literature). This is in contrast to Violante et al., 2023 where 1:3 gives stronger MD than 1:1 does. This may be because Violante et al., 2023 recorded the signal in the direction perpendicular to the hippocampal longitudinal axis in a human cadaver, while we recorded in the radial-in direction in an agar phantom. As expected, random montages induce weaker MD than both literature and optimized-MD montages. We observed a similar pattern in the model (Fig. 12B). From Contact #1 to #3, the model and recordings both show the same relative strength of MD across montages: optMD is the strongest, followed by literature montages (1:1 then 1:3 ratio), a variant of optMD, and then optFocality and random montages. We also note that the model generates about twice higher MD. This may be because the conductivity used in the model (1.0 S/m) is not exactly the conductivity of the agar. This preliminary experimental evidence suggests that individually optimized montages indeed give better stimulation than both literature montages and random montages, highlighting the importance of individualization.

**Fig. 12:**
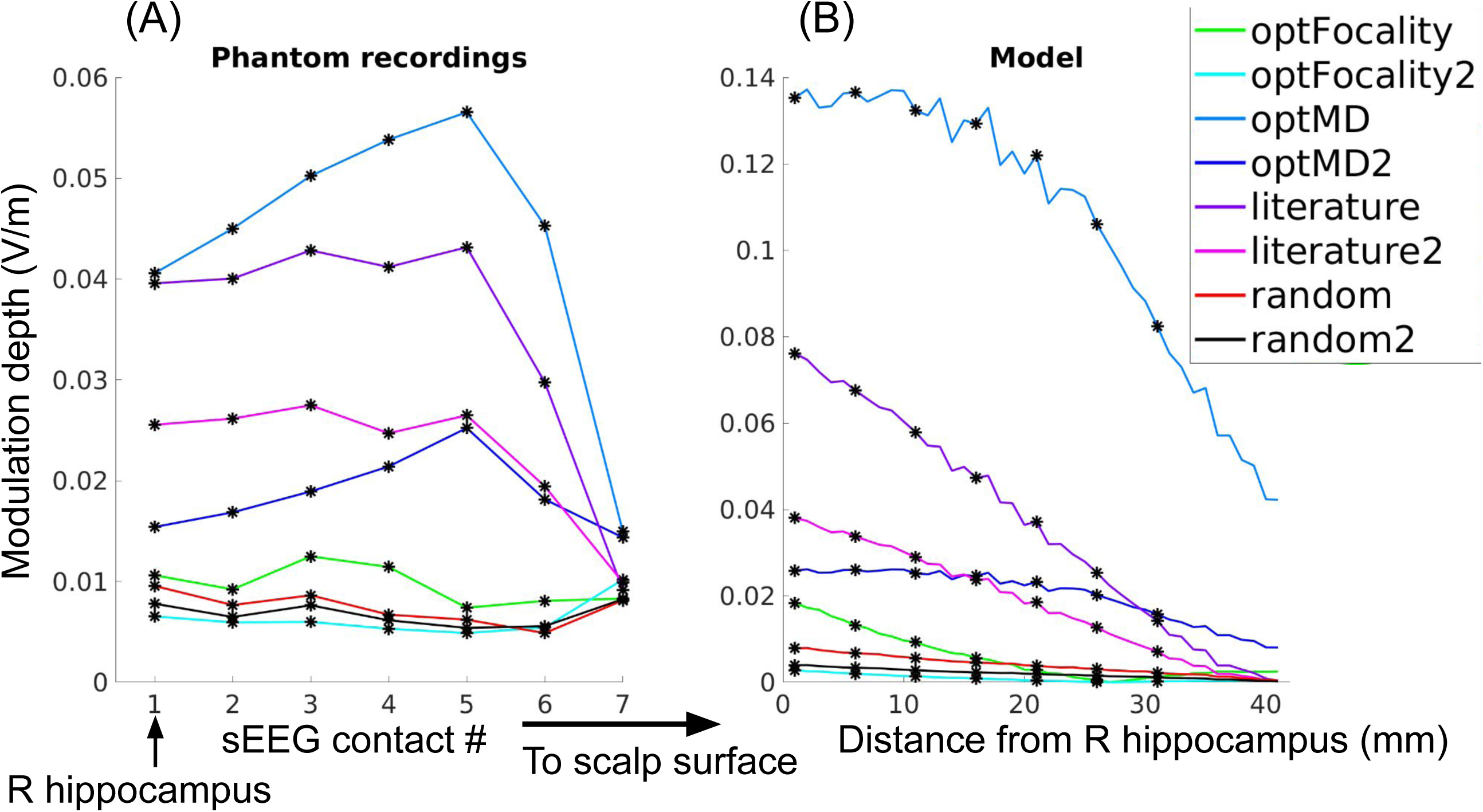
(A) Recorded modulation depth (MD) from the phantom of Subject 1 at the 7 sEEG contacts from the optimized focality (optFocality), optimized MD (optMD), literature (Violante et al., 2023), and random montages. Note no data on the 8th contact as two contacts are needed to compute the electric field. Contact #1 is at the right hippocampus. See Table 2 for details of montages. (B) MD output by the homogeneous-conductivity model of Subject 1, along the line going in radial-out direction from the right hippocampus to the cortical surface. Locations of sEEG contacts are indicated by asteroids. The suffix “2” in the legend means the second montage in each column in Table 2 with current applied at the stimulation electrode altered.

## Discussions

TI has become a popular method for non-invasive brain stimulation recently. However, to the best of our knowledge, all the work studying inter-individual variability used a same (possibly optimized) montage across different individual heads (Acerbo et al., 2022; Hirata et al., 2024; Thiele et al., 2024; von Conta et al., 2022, 2021; Yatsuda et al., 2024; Zhang et al., 2022). This work investigated the inter-individual variability with individually optimized montage and showed *in silico* that there is substantial inter-individual variability in the optimal montages and achieved focality targeting the same brain region. The variability could be due to the differences in head anatomies and tissue conductivities across subjects (Datta et al., 2012; Huang et al., 2017). We found that using a common montage or a montage found in the literature for 2-pair TI will not achieve the maximal focality for each subject at the target. The drop in focality at the target is up to 4 cm when using a common or a literature montage, comparable to the size of some regions of interest in the brain such as the pulvinar, which is a popular region to modulate in schizophrenia (Alemán-Gómez et al., 2020; Kemether et al., 2003). Further sensitivity analysis showed that one can lose focality up to 9 cm when using random montages, for all the three stimulation modalities studied here (2-pair TI, 2-array TI, HD-TES). We also confirmed experimentally the importance of individually optimizing the TI montages in a head phantom.

Therefore, adopting a common montage optimized for one head model targeting a specific region may not guarantee targeted TI stimulation of the same region on other heads, demonstrating the need to individually optimize the montages. This may also help explain why recent work showed that TI fails to induce expected effects when no individual optimization was performed (Iszak et al., 2023).

This work replicated on 25 more heads the previously reported (Huang et al., 2020) superiority of optimized 2-array TI in terms of focality compared to conventional optimized HD-TES on most of the target locations studied. On Target 5, we did not find significant advantage of 2-array TI over HD-TES, probably due to the numerical instability of the optimization algorithm for 2-array TI we noticed in Huang et al., 2020, or specificity of this target location. Ongoing work is developing faster and better algorithms for optimizing 2-array TI so that we can run this optimization across all brain voxels to confirm if the advantage of TI over HD-TES in focality is universal. We also note that the boost in focality from HD-TES to 2-array TI in this work is less than what we previously reported in Huang et al., 2020, as in this work only 72 candidate electrodes were placed on the scalp, excluding those extra 21 electrodes placed at the lower part of the head as in Huang et al., 2020 which may help guide the current to reach the deep brain regions. This may also explain why there is no significant difference between the individually optimized 2-array TI and the common montage on Target 4 which is the deepest in the brain among the six targets studied.

Note that all the head MRIs in the HCP database we used have their faces blurred for protecting the identity of the subjects. However, we did not place any electrodes on the facial areas, and the blurring is not significant enough to affect the structural integrity of the facial areas.

Therefore, we did not expect any significant change in current flow in the model compared to that from using an intact MRI without face blurring. Also, in building all the models in this work we did not utilize the T2-weighted MRIs, as we only see improvement in segmentation from using T2 images in lesioned heads (Huang et al., 2013), while in this work only heads with normal anatomy were modeled.

In this work, we only used one method for optimizing the 2-array TI and a naive search algorithm for the 2-pair TI. Other TI optimization methods exist in the literature, such as using the genetic algorithm to search for the optimal 2 pairs of electrodes (Stoupis and Samaras, 2022), and leveraging modern deep-learning methods to learn the best pattern of the 2 pairs of electrodes (Bahn et al., 2022). There is also a method to search for an optimal montage of multi-pairs of electrodes (Lee et al., 2022). However, we note that all these algorithms take a similar amount of time as ours to solve for one target (in a couple of hours), due to the non-convexness of the TI optimization problem. Deep-learning methods appear to run faster thanks to the ability to run in parallel on a graphic processing unit (GPU). Geng et al., 2025 claims that their algorithm can find the optimal 2-pair solution in 25 seconds. To the best of our knowledge, there is still no algorithm that can solve for the TI optimization in real time, limiting the clinical applications of this technology. Ongoing work is trying to reduce the computational time in solving the TI optimization to make it feasible for practical use.

The initial motivation and advantage of TI over conventional TES is the boosted focality in the deep brain regions (Grossman et al., 2017; Violante et al., 2023). We previously showed that TI without optimization only gives a small increase in focality (Huang and Parra, 2019) and optimization for focality specifically can significantly improve the stimulation focality (Huang et al., 2020). We show the importance of individually optimizing for the focality in this work. Future research should further investigate if increased focality of TI stimulation leads to significantly better functional improvement in areas such as motor and cognitive skills.

This work has some limitations. First, all the results here need to be validated by empirical recordings *in vivo*. Although there are indirect functional *in vivo* imaging on human subjects and electrical recordings on a human cadaver head (Violante et al., 2023), direct *in vivo* recordings of intracranial electrical signals during TI are still lacking in the literature. These data may help determine whether the modulation depth is even the proper quantity to optimize for TI, considering its caveats reported recently (Huang, 2023; Mirzakhalili et al., 2020). We only confirmed the need of individual optimization on a simple head phantom. The results are still preliminary due to the experimental setup where the tube on the phantom is thicker than the sEEG probe, leading to only approximate placement of the probe. This may explain why the model does not exactly match the recordings at all sEEG contacts. Ongoing work on measuring the electrical potential under TI stimulation in a saline-filled tank shows improved focality under 6-pair TI compared to 2-pair TI (unpublished results). Further work is needed to provide empirical evidence supporting the advantage of using multi-channel TI. Secondly, we note that there are feasibility issues implementing 2-array TI in practice, especially when there are overlaps in electrodes across the two frequencies, which may short-cut the current flow and introduce cross-talk across the two frequencies. Ongoing work focuses on improving the algorithm to prevent overlaps of electrodes in the 2 arrays of electrode montages. Also, there is still no device available for 2-array TI. Ongoing work is manufacturing multi-channel (up to 32 per frequency) TI devices for 2-array TI. Previous results on three individual heads showed that one can use as few as 16 electrodes per frequency in 2-array TI to still obtain better focality than that of the 2-pair TI (Huang and Datta, 2021). Thirdly, in this work we did not model other tissue details such as the spongy bone marrow inside the skull, and the anisotropy of white matter. However, our previous study leveraging *in vivo* intracranial electrical recordings on human subjects showed that modeling these details does not provide significant improvement on the TES model accuracy (Huang et al., 2017). Nevertheless, more empirical recordings are needed to test if this is true for TI.

## Acknowledgements

The authors would like to thank Denis Arce, Yishai Valter, and Yaping Huang for their dedicated support and insightful advice in making the head phantom out of agar powder. This work was supported by grants from the National Institute of Health (NIH): 1R44MH126833-01A1 and Department of Education (ED): 91990022C0043.

